## Supplemental Figures 1, 2 & 3 for "Inverted encoding of neural responses to audiovisual stimuli reveals super-additive multisensory enhancement"

**eLife**

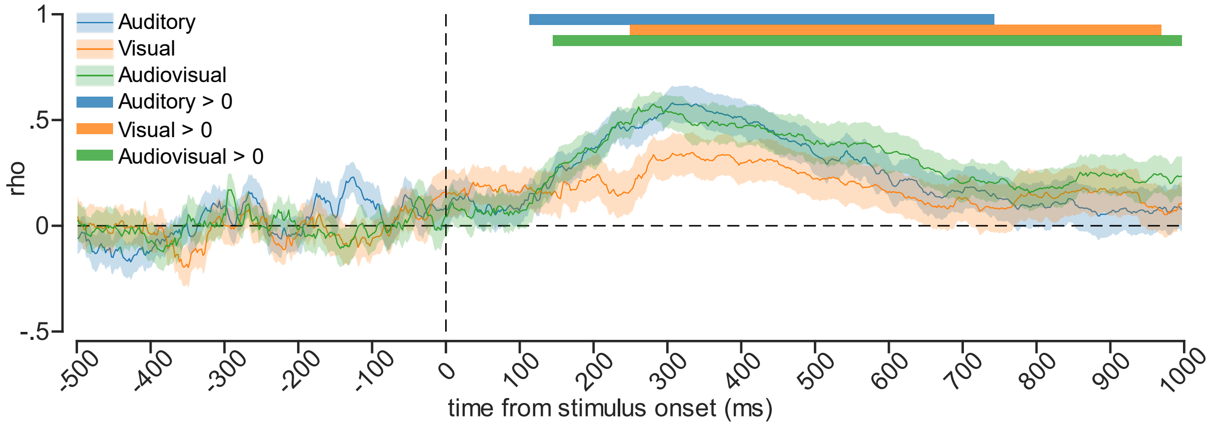


**Figure S1. Correlations reveal a positive relationship between eye position and stimulus position in all conditions.** Correlations (Spearman's rho) between average eye position and stimulus position. Shaded error bars indicate ±1 SEM. Coloured horizontal bars indicate cluster corrected periods that are significantly different from chance (0). Cluster corrected analysis revealed no significant differences between the conditions. If consistent saccades to audiovisual stimuli were responsible for the non-linear decoding shown in Figure 5, we would expect a greater correlation between horizontal eye-position and stimulus location for the audiovisual condition compared with the unisensory conditions.

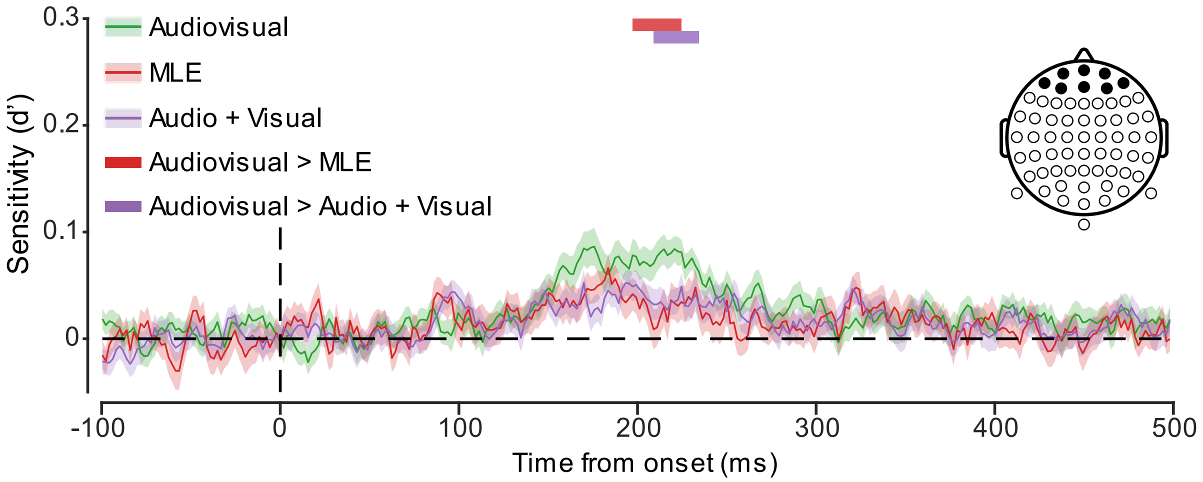


**Figure S2. Decoding sensitivity from frontal electrodes.** Predicted (optimal sensitivity through MLE and aggregate A+V) and actual audiovisual decoding sensitivity for decoding stimulus locations from fontal neural activity. Shaded error bars indicated ±1 SEM. Sensitivity is markedly lower than that from posterior neural activity (Figure 5), suggesting eye movements are not a primary contributor to non-linear multisensory enhancements.

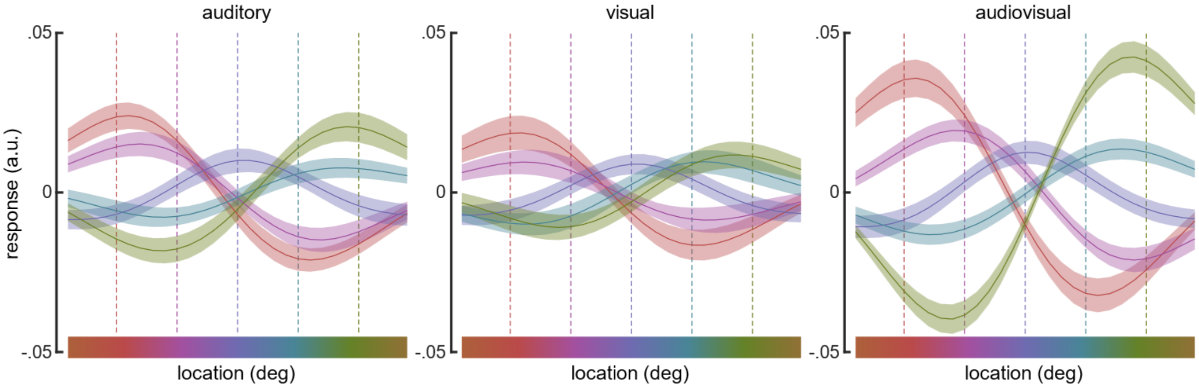


**Figure S3. Channel activity.** To represent the channel activity of the forward model, we reconstructed the tuning curves from each of the five channel responses for each location (±[0, 7.5, 15]º). They were computed by taking the dot product of the channel responses and the forward model. Vertical dashed lines indicate the stimulus positions. Channel responses are colour coded, and shaded regions indicate ±1 SEM.
